# Cerebellar Modulation of Mesolimbic Dopamine Transmission is Functionally Asymmetrical

**DOI:** 10.1101/615815

**Authors:** Zade R. Holloway, Nick B. Paige, Josiah F. Comstock, Hunter G. Nolen, Helen J. Sable, Deranda B. Lester

## Abstract

Cerebral and cerebellar hemispheres are known to be asymmetrical in structure and function, and previous literature supports that asymmetry extends to the neural dopamine systems. Using *in vivo* fixed-potential amperometry with carbon-fiber microelectrodes in anesthetized mice, the current study assessed hemispheric lateralization of stimulation-evoked dopamine in the nucleus accumbens (NAc) and the influence of the cerebellum in regulating this reward-associated pathway. Our results suggest that cerebellar output can modulate mesolimbic dopamine transmission, and this modulation contributes to asymmetrically lateralized dopamine release. Dopamine release did not differ between hemispheres when evoked by medial forebrain bundle (MFB) stimulation; however, dopamine release was significantly greater in the right NAc relative to the left when evoked by electrical stimulation of the cerebellar dentate nucleus (DN). Furthermore, cross-hemispheric talk between the left and right cerebellar DN does not seem to influence mesolimbic release given that lidocaine infused into the DN opposite to the stimulated DN did not alter release. These studies may provide a neurochemical mechanism for studies identifying the cerebellum as a relevant node for reward, motivational behavior, saliency, and inhibitory control. An increased understanding of the lateralization of dopaminergic systems may reveal novel targets for pharmacological interventions in neuropathology of the cerebellum and extending projections.

## Introduction

No longer considered a structure primarily for motor coordination, the cerebellum is now known to contain three distinct regions that contribute to sensorimotor, limbic, and cognitive processes [1]. Cerebellar and cerebral systems work in concert to sharpen the timing of these neural operations [2,3], and each cerebellar hemisphere is connected to multiple closed-loop cortical neural networks in the contralateral cerebral hemispheres, providing an anatomical basis for a cerebellar role in cognition [4–6] and a cerebellar mirroring of functional specializations in the cerebrum [7]. Specifically, the cerebellum receives input from the cerebral hemispheres via pontine nuclei in the brainstem, and relays to the contralateral cerebral cortex via cerebellar Purkinje cells and their projections to the dentate nucleus (DN), which provides the sole output from the cerebellum to the cerebrum [8–10].

Cerebrocerebellar networks have been shown to be asymmetrical in structure and function in many species including birds, rodents, primates, and humans [11–16]. Clinical and preclinical studies support the association of the left cerebral hemisphere with communication functions and the right cerebral hemisphere with spatial reasoning [17,18]. Due to contralateral connections between cerebrocerebellar systems, the cerebellar hemispheres parallel these specializations. Imaging and lesion studies in humans have found the left cerebellar hemisphere to be involved in visuo-spatial operations [19–22], and a right cerebellar involvement in language processes [23–25].

Likely related to these behaviorally-associated asymmetries, the bilateral hemispheres of the brain also contain lateralized neurotransmitter systems in cortical and subcortical regions, and certain experiences have been shown to enhance this lateralization. For example, rats that were handled in their early life showed a significant left/right asymmetry (R>L) in dopamine levels in the NAc [11]. Other studies in rats show greater concentrations of DOPAC/DA in the right cortex and nucleus accumbens in comparison to the same systems in the left hemisphere [26]. Increased dopamine levels in the right prefrontal cortex of adult rats were found to be strongly correlated with anxiety responses in the elevated plus-maze test [27], and dopamine receptor blockade in the right mPFC but not the left mPFC of handled rats resulted in elevated stress and hormone levels [28]. These studies provide evidence of hemispheric imbalance in the mesocorticolimbic dopamine system.

Many neurophysiological disorders are characterized by altered profiles of mesocorticolimbic dopaminergic transmission, such as addiction, ADHD, and schizophrenia [29–31], and interestingly, cerebellar pathology and specifically Purkinje cell dysfunction are being considered as substrates in these and other psychiatric disorders [32–34]. A direct pathway from the deep cerebellar nuclei to the VTA has been identified [35]. Mittleman and colleagues [36] used cerebellar DN electrical stimulation to evoke dopamine efflux in the medial prefrontal cortex (mPFC) of Lurcher mutant mice, a common model of autism spectrum disorder with 100% loss of Purkinje cells within the first 4 weeks of life. The Lurcher mutants exhibited attenuation in DN-evoked mPFC dopamine release compared to controls, suggesting that developmental loss of Purkinje cells in the cerebellum, similar to that of autism spectrum disorder, can lead to a disruption in mPFC dopamine transmission. In further studies, these researchers found reorganization of cerebello-cortico circuitry in Lurcher mutants and Fmr1 mice, another genetic model that exhibits dysfunction or absence of Purkinje cells in the cerebellum [37]. The reorganization of the DN to mPFC pathways included altered relative influence of the ventral tegmental area (VTA) and thalamic nuclei, with the mutant mice showing a stronger dependence on thalamic nuclei compared to control mice. This shift in cerebellar modulation towards the ventral lateral thalamus and away from the VTA leads to speculation about the cerebellum’s influence on dopaminergic functioning not only in the mPFC but also the nucleus accumbens (NAc).

Neural fibers between the VTA and NAc constitute one of the most densely innervated dopamine pathways in the brain [38,39]. Dopamine release in the NAc is known to be associated with reward and motivational processes [40–43], and disruption to normal dopamine processing, including hemispheric balance, can lead to a host of motor and cognitive deficits. For example, decreased motivation and novelty-seeking often observed in patients with Parkinson’s disease are related to asymmetry of dopamine [44] and individual differences in incentive motivation or sensitivity to natural rewards in humans has also been associated with increased asymmetry in dopaminergic systems [45].

Using *in vivo* fixed potential amperometry in anesthetized mice, Experiment 1 of the present study aimed to distinguish any asymmetries between the mesolimbic dopamine pathways by stimulating the medial forebrain bundle (MFB), which consists of the dopaminergic axons projecting from the VTA to NAc, and recording dopamine release in the NAc in each hemisphere. In Experiment 2, we assessed cerebellar influence of NAc dopamine lateralization by comparing DN stimulation-evoked dopamine release in both hemispheres. The DN has contralateral glutamatergic projections to reticulotegmental nuclei that, in turn, project to pedunculopontine nuclei (PPT), which projects to and stimulates dopamine cell bodies in the VTA [46,47]. For this reason, the present study stimulated the DN located contralateral to the NAc recording site (left DN stimulation with right NAc recording and vice versa). Lastly, in Experiment 3, we examined the potential cross-hemispheric influence of cerebellar DN on this dopaminergic pathway. During contralateral DN stimulation-evoked dopamine recordings, separate groups of mice received an infusion of either lidocaine or phosphate-buffered saline (PBS; vehicle control) into the ipsilateral DN to determine if communication between each cerebellar DN can influence the contralateral, active pathway being stimulated. (See Figure 1 for a depiction of the experimental configurations.) These experiments are the first to systematically examine the contribution of the cerebellum on lateralized stimulation-evoked phasic dopamine release in the NAc. An improved understanding of cerebello-cortico circuitry may help identify targets for pharmacological interventions in neuropathologies related to dopamine dysfunction.

**Figure 1.**
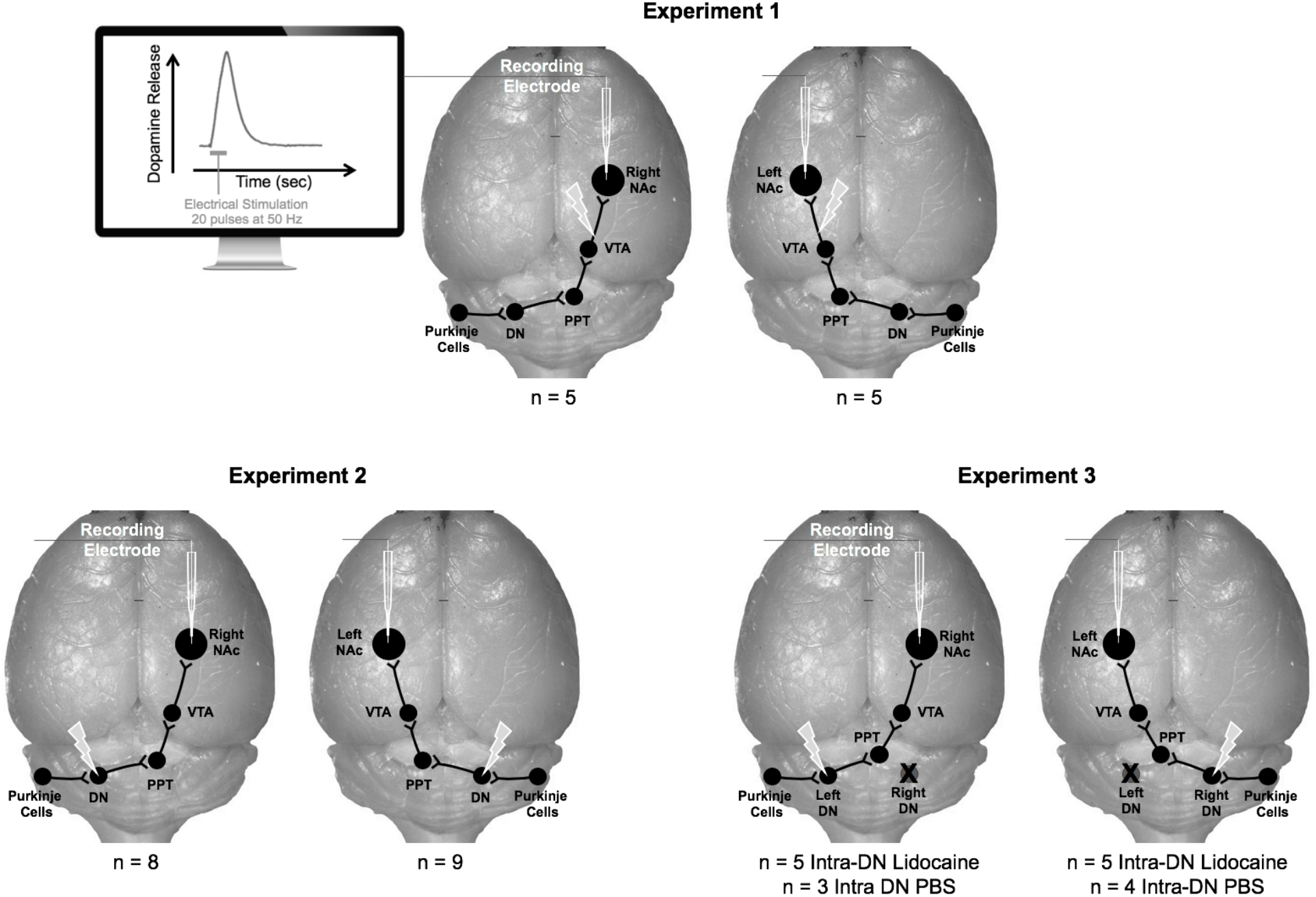
Depiction of Experimental Design. In experiment 1, stimulating electrodes were positioned in the right or left medial forebrain bundle, and carbon fiber recording electrodes were positioned in the ipsilateral nucleus accumbens (NAc). In experiment 2, stimulating electrodes were positioned in the right or left cerebellum dentate nucleus (DN), and recording electrodes were positioned in the contralateral NAc. Experiment 3 consisted of the same stimulating and recording positions as experiment 2, with an added infusion cannula for lidocaine or phosphate buffered saline (PBS) in the DN contralateral to the stimulation. Pedunculopontine tegmentum = PPT, ventral tegmental area = VTA.

## Materials and Methods

### Animals

Twenty-seven male C57BL/6J mice (Jackson Laboratories, ME) were housed 3-5 per cage in a temperature-controlled environment (21±1 °C) on a 12 hr light/dark cycle with (lights on at 0600) and given food and water available ad libitum. All experiments were approved by the Institutional Animal Care and Use Committee at the University of Memphis and conducted in accordance with the National Institutes of Health Guidelines for the Care and Use of Laboratory Animals. In order to maintain the principle of reduction related to scientific experiments on animals [48], efforts were made to minimize the number of mice used. Sample sizes were determined based on G*Power analysis [49] and effect sizes from our previous amperometric results [50–52]. Efforts were also made to minimize pain and discomfort.

### Surgery

Mice were anesthetized with urethane (1.5 g/kg, i.p.) and placed in a stereotaxic frame (David Kopf Instruments, Tujundga, CA) ensuring a flat skull. Body temperature was maintained at 36±0.5 °C with a temperature-regulated heating pad (TC-1000; CWE, NY). All stereotaxic coordinates are in mm from bregma, midline, and dura according to the mouse atlas of Paxinos and Franklin [53]. In each mouse, a concentric bipolar stimulating electrode (SNE-100, Rhodes Medical, CA) was implanted into either the left cerebellar DN (AP −6.25, ML +2.0, and DV −2.0), or right DN (AP −6.25, ML −2.0, and DV −2.0), or either the left MFB (AP −2.0, ML +1.1, DV −4.0) or right MFB (AP −2.0, ML −1.1, DV −4.0). A stainless-steel auxiliary and Ag/AgCl reference electrode combination was placed on the surface of cortical tissue contralateral to the stimulation electrode and −2.0 mm from bregma, and a carbon fiber recording electrode was positioned in either the left NAc (AP +1.5, ML +1.0, and DV −4.0) or the right NAc (AP +1.5, ML −1.0, and DV −4.0). For MFB stimulations, the recording electrode was placed in the ipsilateral NAc; however, due to contralateral connections in cerebrocerebellar circuity, the recording electrode was placed contralateral to cerebellar DN stimulation [46,47]. Pharmacological studies from our lab have confirmed the recorded current changes in the NAc to be dopamine dependent [50,51].

### Dopamine Recordings and Drug Infusions

Fixed potential amperometry, also known as continuous amperometry, coupled with carbon fiber recording microelectrodes has been confirmed as a valid technique for real-time monitoring of stimulation-evoked dopamine release [52,54,55]. All amperometric recordings were made within a Faraday cage to increase signal to noise ratio. A fixed potential (+0.8 V) was applied to the recording electrode, and oxidation current was monitored continuously (10 K samples/sec) with an electrometer (ED401 e-corder 401 and EA162 Picostat, eDAQ Inc., Colorado Springs, CO) filtered at 50 Hz. A series of cathodal current pulses was delivered to the stimulating electrode via an optical isolator and programmable pulse generator (Iso-Flex/Master-8; AMPI, Jerusalem, Israel). The stimulation protocol consisted of 20 monophasic 0.5 ms duration pulses (800 μAmps) at 50 Hz every 60 seconds to establish a baseline dopamine response. MFB and DN stimulation-evoked dopamine was monitored for 30 minutes in each mouse. Following these baseline recordings, a random subset of mice received a 1.0 μL infusion (over 1.0 min) of either PBS (vehicle control) or the local anesthetic lidocaine (4%) into the DN contralateral to the stimulation site, and dopamine recordings continued for 30 min. Lidocaine blocks sodium channels and has been used during amperometric dopamine recordings to temporarily block functioning in a local brain site with peak lidocaine responses occurring between 2-5 min post infusion [52]. At the conclusion of the amperometric recordings, recording electrodes were calibrated in vitro with dopamine solutions (0.2-1.2 μM) administered by a flow injection system [56,57]. Thus, change in dopamine oxidation current (nAmp) was converted to dopamine concentration (μM).

### Histology

Upon the completion of each amperometric recording, an iron deposit was made in the stimulation site by passing direct anodic current (100 μA and 1 mA, respectively) for 10 sec through the stimulating electrodes, and 1.0 μL cresyl violet stain was infused into the cannula site. Mice were euthanized with a 0.25 ml intracardial injection of urethane (0.345 g/ml). Brains were removed, immersed in 10% buffered formalin containing 0.1% potassium ferricyanide (which causes a redox reaction at the stimulation site resulting in a Prussian blue spot), and then stored in 30% sucrose/10% formalin solution for at least 1 week prior to sectioning. Using a cryostat at −20°C, 30 μm coronal sections were sliced, and electrode placements were determined under a light microscope and recorded on representative coronal diagrams confirming the intended sites were stimulated [53].

### Drugs

Urethane (U2500), lidocaine, (L7757), and dopamine (H8502) were obtained from Sigma-Aldrich Chemical (St Louis, MO). Urethane was dissolved in distilled water, and lidocaine and dopamine were dissolved in PBS (pH 7.4).

### Data Analysis

To quantify MFB and DN stimulation-evoked dopamine efflux, pre-stimulation current values were normalized to zero, and data points occurring 0.25 sec pre- and 50 sec post-onset of the stimulation were extracted. Dopamine release was quantified as the magnitude of the response (peak minus baseline). Independent samples t-tests were used to assess hemispheric differences in NAc dopamine release. Stimulation-evoked dopamine release was also measured 5 min following intra-DN infusion. A twoway mixed ANOVA was used to determine the effect of drug infusion (PBS or lidocaine) and time (pre- or post-infusion) on dopamine release. Dopamine release post-infusion was also converted to percent change (with the pre-infusion concentration being 100%), and an independent samples *t*-test was used to determine if there was a significant difference between PBS and lidocaine.

## Results

### Histology

The tips of the stimulating electrodes were positioned within the anatomical boundaries of the DN or MFB. Figure 2A-B depicts the medial/lateral and dorsal/ventral placement ranges for the DN and MFB, respectively [53]. The anterior/posterior positioning ranges for the stimulating electrodes were −6.00 to −6.36 mm from bregma for the DN and −1.82 to −2.18 mm from bregma for the MFB. The active portions of the carbon fiber recording electrodes were positioned within the anatomical boundaries of the NAc core (Figure 2C) [53], with the anterior/posterior positioning ranges being +1.34 to +1.70 from bregma.

**Figure 2.**
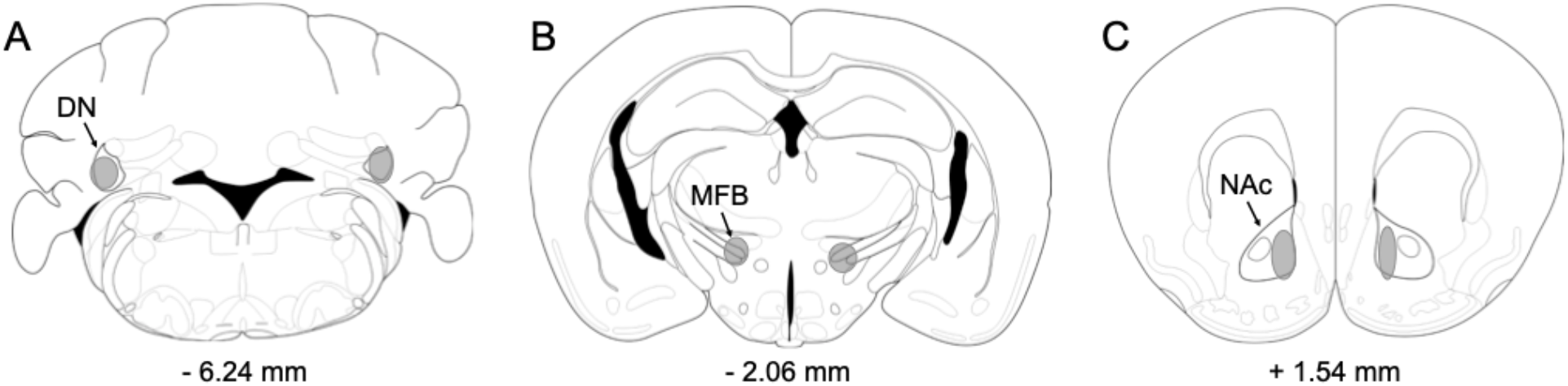
Representative coronal sections of the mouse brain (adapted from the atlas of Paxinos & Franklin, 2001), with gray-shaded areas indicating the placements of stimulating electrodes in the A) cerebellar dentate nucleus (DN) or B) medial forebrain bundle (MFB) and amperometric recording electrodes in the C) nucleus accumbens (NAc). Numbers correspond to mm from bregma.

### Experiment 1: NAc dopamine release following ipsilateral stimulation of the MFB

NAc dopamine release was quantified in each hemisphere as a function of peak height following electrical stimulation of the ipsilateral MFB. No differences were observed between the MFB stimulation-evoked dopamine release in the right NAc (M ± SEM: 1.614 uM ± 0.466, n = 5) compared to the left NAc (1.513 uM ± 0.357, n = 5); *t* (8) = −0.172, *p* = .87 (Fig. 3).

**Figure 3.**
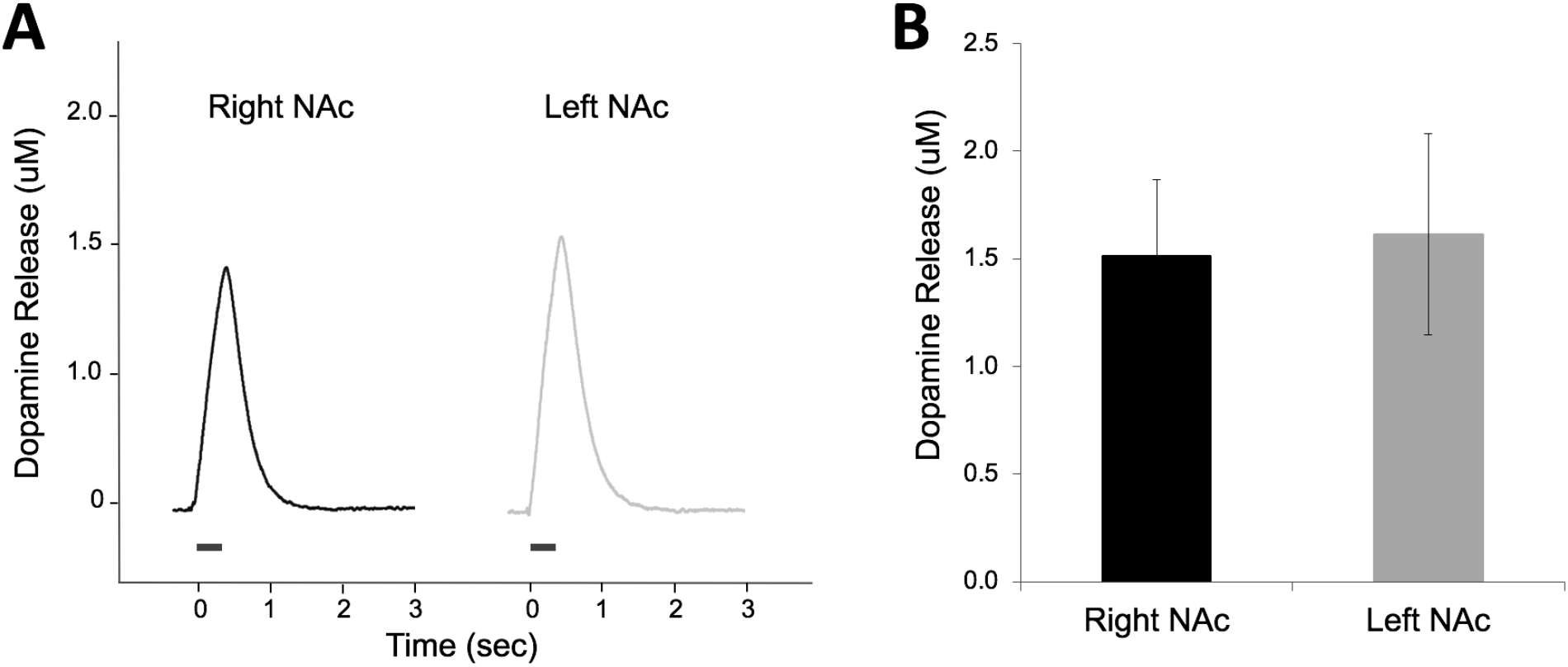
Amperometric recordings of dopamine release in the right or left nucleus accumbens (NAc) in response to electrical stimulation of the ipsilateral medial forebrain bundle. (A) Profiles illustrate example responses from each recording site. Time zero indicates the start of the train of 20 pulses at 50 Hz. (B) No mean (± SEM) differences in dopamine release were observed between hemispheres.

### Experiment 2: NAc dopamine release following contralateral stimulation of the cerebellar DN

NAc dopamine release was quantified in each hemisphere as a function of peak height following electrical stimulation of the contralateral DN. DN stimulation-evoked dopamine release was significantly greater in the right NAc (M ± SEM: 0.018 uM ± 0.002) compared to the left NAc (0.011 uM ± 0.001); *t* (15) = −3.47, *p* < .01, *d* = 1.67 (Fig. 4).

**Figure 4.**
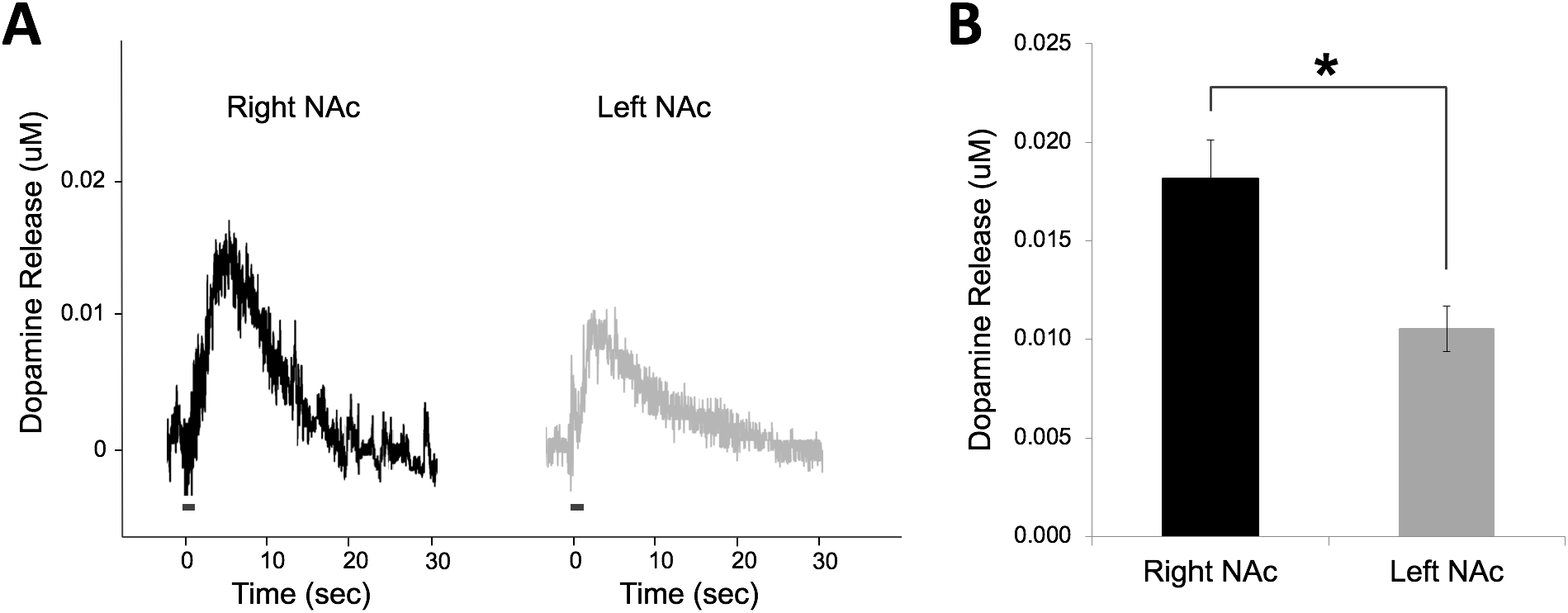
Amperometric recordings of dopamine release in the right or left nucleus accumbens (NAc) in response to electrical stimulation of the contralateral dentate nucleus (DN) of the cerebellum. (A) Profiles illustrate example responses from each recording site. Time zero indicates the start of the train of 20 pulses at 50 Hz. (B) Mean (± SEM) differences in dopamine release were observed between hemispheres. * indicates *p* < .01.

### Experiment 3: NAc dopamine release following deactivation of the ipsilateral cerebellar DN

During contralateral DN stimulation-evoked dopamine recordings, separate mice received an infusion of either PBS or lidocaine into the ipsilateral DN to determine the impact of hemispheric DN interactions on mesolimbic dopamine transmission. In the right NAc (electrical stimulation in the left DN and infusion into the right DN), a two-way mixed ANOVA revealed no significant interaction between the infusion (PBS or lidocaine) and time (pre or post-infusion) on dopamine release, *F*(1, 6) = 0.13, *p* = .73, and no main effect of infusion on dopamine release, *F*(1, 6) = 0.01, *p* = .91. Similarly, in the left NAc (electrical stimulation in the right DN and infusion into the left DN), a twoway mixed ANOVA revealed no significant interaction between the infusion (PBS or lidocaine) and time (pre- or post-infusion) on dopamine release, *F*(1, 7) = 0.39, *p* = .55, and no main effect of infusion on dopamine release, *F*(1, 7) = 0.36, *p* = .57. These results indicate that in both hemispheric NAc recordings, dopamine release was not altered by either infusion (PBS or lidocaine), suggesting DN cross-talk is not significantly influencing NAc dopamine release. Figure 5 shows this data in terms of percent change with dopamine recordings prior to infusion being 100%. Correspondingly, no differences in percent change in dopamine release were observed between lidocaine and PBS infusions in either the left NAc recordings [*t* (7) = 0.33, *p* = .76, Fig. 5A] or right NAc recordings [*t* (6) = −0.61, *p* = .57, Fig. 5B].

**Figure 5.**
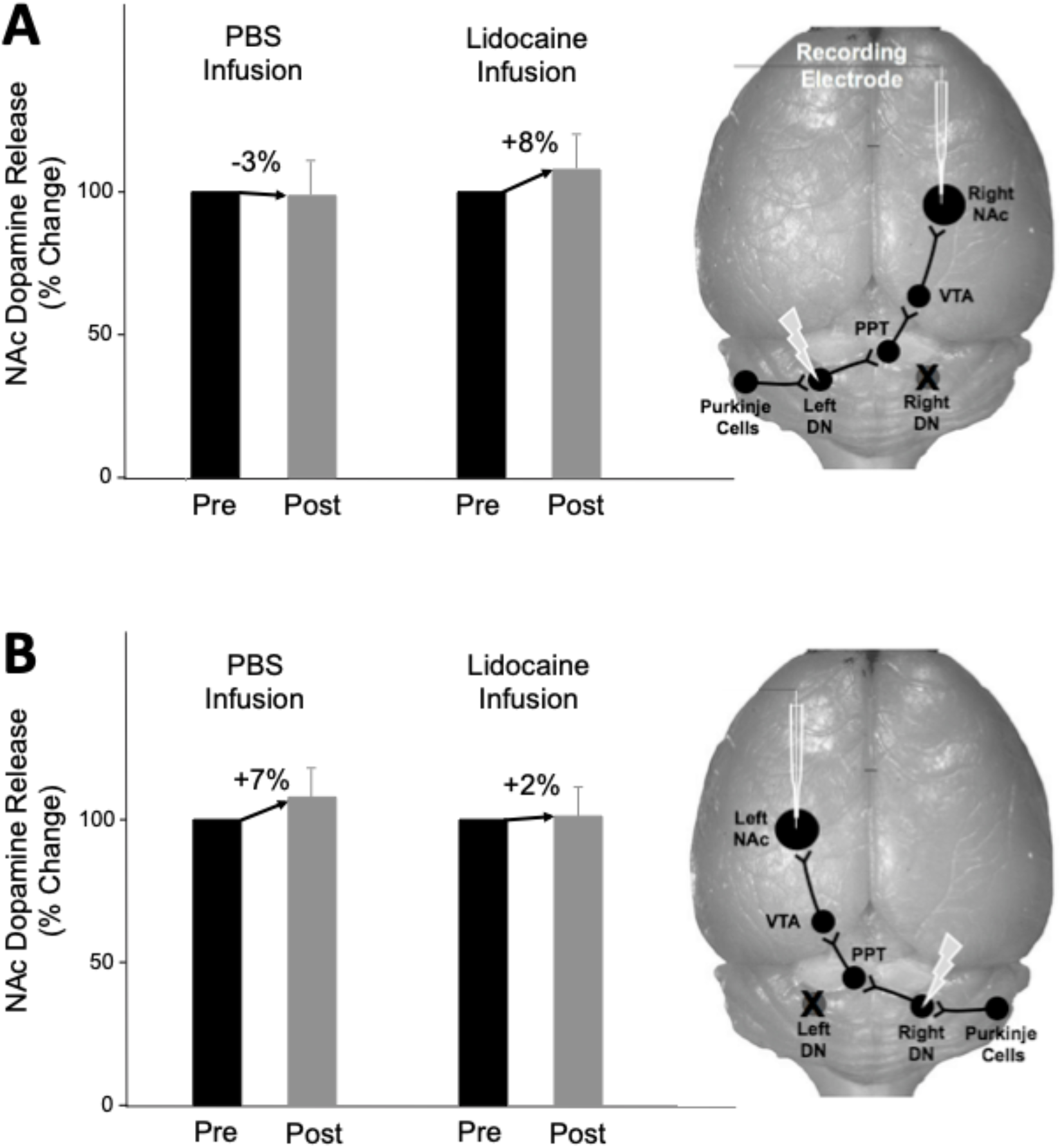
Mean (± SEM) nucleus accumbens (NAc) dopamine release in response to electrical stimulation of the contralateral cerebellar dentate nucleus (DN) pre and post infusion of PBS or lidocaine in the ipsilateral DN. Neither PBS of lidocaine infusion significantly altered dopamine release in the NAc (A: right NAc recordings, B: left NAc recordings).

## Discussion

The current study assessed the hemispheric lateralization of stimulation-evoked dopamine in the NAc and the influence of the cerebellum in regulating this reward-associated pathway. Results from Experiment 1 show that the mesolimbic pathway itself is not responsible for asymmetrical lateralization of dopamine release, given that NAc dopamine release did not differ between hemispheres when evoked by ipsilateral MFB stimulation. The mesolimbic dopamine pathways are known to be ipsilaterally dominated, with less than 10% of fibers from the VTA crossing hemispheres to reach the contralateral NAc [58], and our findings suggest that these parallel pathways act functionally similar when equally stimulated. Instead, dopaminergic asymmetry may originate, at least in part, from the influence of the cerebellum on these pathways. In Experiment 2, we show that stimulation of the cerebellar DN can elicit NAc dopamine release in the contralateral hemisphere, and this dopamine release was significantly greater in the right NAc relative to the left.

Furthermore, cross-hemispheric talk between the right and left cerebellar DN does not seem to influence mesolimbic dopamine release given that, in Experiment 3, when lidocaine was infused into the DN opposite the electrically stimulated DN to inactive this system, dopamine release was not altered. We have previously shown that this infusion protocol and lidocaine dose can deactivate local neural activity from 2 to 20 min post infusion [52]. The present findings exhibiting no cross-hemispheric talk between the dentate nuclei is not surprising given that these structures are physically separated by the cerebellar vermis [59]. Furthermore, the cerebro-cerebellar networks are thought to be laterally independent circuits, with cortical hemispheric connections primarily running through the corpus callosum [60–62].

The MFB-evoked dopamine release profiles in the NAc observed in the present study are consistent with those previously published [51,52,63]. When comparing dopamine release in the NAc, mPFC, and amygdala, we have previously characterized NAc dopamine release to be higher in concentration and quickly cleared from the synapse, indicating greater synaptic confinement, relative to the other brain regions [51]; however, in the present study, NAc dopamine release elicited by cerebellar DN stimulation was attenuated in concentration and slower to clear from the synapse, allowing for greater diffusion beyond the synaptic release site, relative to MFB stimulated dopamine release. The profile of DN-evoked dopamine release in the NAc more closely fits the description of volume transmission. In contrast to point-to-point synaptic contacts, volume transmission provides a communication mode that is temporally slower, broader in anatomical reach, and more suited to modulatory/tuning functions [64]. Thus, the present findings demonstrate a functional regulatory role of the cerebellum over mesolimbic dopamine activity and provide a neurochemical mechanism for studies identifying the cerebellum as a relevant node for reward, motivational behavior, saliency, and inhibitory control [65–68].

Cerebral and cerebellar hemispheres are known to be asymmetrical in structure and function [12,13], and studies are mounting to show that this asymmetry extends to the mesolimbic dopamine system. Although dopaminergic asymmetries are not consistently documented in the literature and seem to vary based on age, gender, species, and strain [for review see 69], numerous studies using methods of protein analyses in rodents have shown hemispheric distributions similar to the present results, greater levels of dopamine and its metabolites in the right NAc relative to the left [11,26,70,71]. It has been suggested that individual differences to natural rewards are a product of asymmetry in dopamine systems [72]. The present results support the notion that reward processes in the brain may be lateralized between cerebello-cortico circuitry, which has considerable applications for disorders involving dysfunction of the subcortical dopamine functioning, disorders such as schizophrenia, Parkinson’s, ADHD, and addiction [29–31]. For example, patients with unlilateral onset of Parkinon’s disease often develop an asymmetry of dopamine deficiency [73,74], with behavioral deficits not limited to motor functioning. In one study, patients whose motor symptoms began on the left side of the body performed more poorly on cognitive tests than those with right-side onset, leading to the conclusion that damage to right-hemisphere dopamine plays a greater role in associated cognitive decline than left-hemisphere depletion [75]. In the future, treatment for such symptoms may be optimal with if applied differentially in each hemisphere [72].

Cerebellar-mediated dopamine pathways have previously been shown to exhibit plasticity and compensatory changes in the neural circuitry of rodent models of autism, providing a foundation for the cerebellum to develop unique connections between cerebral hemispheres [67]. Some hypothesize that the corpus callosum enables hemispheric specializations, allowing one hemisphere to reconfigure circuits and adapt to certain environmental changes while the other hemisphere preserves existing functions [77]. Others have suggested that lateralization may occur though the action of steroid hormones [78]. Despite varying perspectives on mechanism of action or developmental precursors, neural circuits specialize their connections to use resources more efficiently and minimize wiring [62]. It appears that this specialized processing is represented in the brain as analogue signals, namely changes in the concentration of messenger molecules in the synaptic space [79]. The differences observed in concentrations of dopamine release in left and right cerebellar-NAc circuits may provide a foundation for the divergent types of information processed and transmitted between the reward circuits of each hemisphere. Although each hemisphere may contain homologous neural substrates, lateralized dopamine release patterns allow for anatomically defined circuitry to be repurposed and used for other adaptive behaviors [80]. Further examination into the functional relationship between lateralized cerebrocerebellar networks may help stimulate new insight to understanding hemispheric specializations in neurodevelopment and lead to novel targets for pharmacological interventions.

